# Tafazzin deficiency in mouse mesenchymal stem cells potentiates their immunosuppression and impairs activated B lymphocyte immune function

**DOI:** 10.1101/2021.09.07.459330

**Authors:** Hana M. Zegallai, Ejlal Abu-El-Rub, Folayemi Olayinka-Adefemi, Laura K. Cole, Genevieve C. Sparagna, Aaron J. Marshall, Grant M. Hatch

## Abstract

Barth Syndrome (BTHS) is a rare X-linked genetic disorder caused by mutation in the TAFAZZIN gene which encodes the cardiolipin (CL) transacylase tafazzin (Taz). Taz deficiency in BTHS patients results in reduced CL in their tissues and a neutropenia which contributes to the risk of infections. However, the impact of Taz deficiency in other cells of the immune system is poorly understood. Mesenchymal stem cells (MSCs) are well known for their immune inhibitory function. We examined whether Taz-deficiency in murine MSCs impacted their ability to modulate lipopolysaccharide (LPS)-activated wild type (WT) murine B lymphocytes. MSCs from tafazzin knockdown (TazKD) mice exhibited a 50% reduction in CL compared to wild type (WT) MSCs. However, mitochondrial oxygen consumption rate and membrane potential were unaltered. In contrast, TazKD MSCs exhibited increased glycolysis compared to WT MSCs and this was associated with elevated proliferation, phosphatidylinositol-3-kinase expression and expression of the immunosuppressive markers indoleamine-2,3-dioxygenase, cytotoxic T-lymphocyte-associated protein 4, interleukin-10, and cluster of differentiation 59. When co-cultured with LPS-activated WT B cells, TazKD MSCs inhibited B cell proliferation and growth rate and reduced B cell secretion of IgM to a greater extent than B cells co-cultured with WT MSCs. In addition, co-culture of LPS-activated WT B cells with TazKD MSCs induced B cell differentiation toward potent immunosuppressive phenotypes including interleukin-10 secreting plasma cells and B regulatory cells compared to activated B cells co-cultured with WT MSCs. These results indicate that Taz deficiency in MSCs enhances MSCs-mediated immunosuppression of activated B lymphocytes.

## Introduction

Barth syndrome (BTHS) is a rare X-linked genetic disorder caused by mutations in the TAFAZZIN gene localized on chromosome Xq28 (Clarke et al., 2013; Hauff and Hatch, 2006). The product of the TAFAZZIN gene is the tafazzin (Taz) protein, a phospholipid transacylase enzyme that plays a critical role in maturation of the inner mitochondrial membrane phospholipid cardiolipin (CL) and remodels nascent CL to mature CL (Xu et al., 2006). CL comprises approximately 15-20% of the inner mitochondrial membrane phospholipid mass and is required for the activation of many enzymes and proteins involved in mitochondrial function (Osman et al., 2011; Zegallai and Hatch, 2021). BTHS patients exhibit CL deficiency and abnormal CL acyl chain composition leading to impairment and inhibition of ATP synthesis from oxidative phosphorylation (Ikon and Ryan, 2017). BTHS is characterized by cardiomyopathy, skeletal myopathy, neutropenia, and growth retardation (Dudek and Maack, 2017). Cardiomyopathy is the main cause of mortality in BTHS patients. However, BTHS patients are vulnerable to severe infections as approximately 70% of patients develop neutropenia (Clarke et al., 2013). The mechanism for the neutropenia in BTHS is not fully understood. In addition, how other cells of the immune system might contribute to immunodeficiency in BTHS patients remains unclear.

Preclinical and clinical studies indicate that mesenchymal stem cells (MSCs) are among the best stem cells for treating many degenerative and chronic disorders (Shammaa et al., 2020). They have gained widespread attention due to their broad therapeutic applications and can be easily isolated from many sources including bone marrow, adipose tissues, umbilical cord and others (Klingemann et al., 2008). MSCs modulate and regulate many cells of the immune system through modification of their activity and differentiation portal (Kaundal et al., 2018). B cells are crucial for humoral immunity of the adaptive immune system (LeBien and Tedder, 2008). Altered immune function of B cells is observed in autoimmune diseases where B cell activation is uncontrolled and exaggerated or in immunodeficiency diseases where B cells are poorly activated and recruited (Lin and Lu, 2020; Moise et al., 2010). The recruitment, activation, differentiation and maturation of B cells is a complicated process and require tight control. Activated B cells secrete antibodies, such as immunoglobulin-M (IgM) and immunoglobulin-G (IgG), and cytokines, such as interleukin-10 (IL-10) (Vazquez et al., 2015). The presence of MSCs at the site of B cell activation and maturation modulate B cell function and differentiation. MSCs prevent B cell proliferation and promote their arrest at the G0-G1 phase of the cell cycle and they secrete a cocktail of cytokines, including IL-10, transforming growth factor-β, prostaglandin E_2_ and indoleamine 2,3, dioxygenase (IDO), which inhibit B cell terminal differentiation (Liu et al., 2020a; Ma and Chan, 2016). In addition, MSCs promote differentiation of B cells into B regulatory cells (Bregs) that induce an immune-tolerance state and impede differentiation of B cells into plasma cells and as a result suppress immunoglobulin secretion (Liu et al., 2020a). Moreover, MSCs interaction with B cells trigger the release of the costimulatory immunoglobulin programmed death ligand-1 which attenuates B cell immune function (Liu et al., 2020b). Thus, the immunosuppressive properties of MSCs are important in regulating and tightly controlling the function of B cells of the immune system. However, the immune-inhibitory role of MSCs can be modulated and enhanced by genetic modifications which may affect the function of other immune cells and lead to an immunodeficient state. To date no studies have examined how Taz deficiency in MSCs impacts the function of other cells of the immune system. In the current study, we demonstrate for the first time that Taz deficiency in murine MSCs attenuate activated murine B cell function. Our results provide novel mechanistic information on the role of Taz in regulating immune system function.

## Materials and Methods

### Experimental animals

All experimental procedures in this study were performed with approval of the University of Manitoba Animal Policy and Welfare Committee in accordance with the Canadian Council on Animal Care guidelines and regulations. This study is reported in accordance with ARRIVE guidelines. All animals were housed in pathogen-free facility (12 h light/dark cycle) at the University of Manitoba. Taz knockdown (TazKD) mice were generated by mating transgenic male mice containing doxycycline (dox) inducible Taz specific short-hair-pin RNA (shRNA) (B6.Cg-Gt(ROSA)26Sortm1(H1/tetO-RNAi:Taz,CAG-tetR)Bsf/ZkhuJwith female C57BL/6J mice from Jackson Laboratory (Boston, MA). Taz knock down was maintained postnatally by administering dox (625 mg of dox/kg of chow) as part of the rodent chow (Rodent diet TD.01306 from Harlan, Indianapolis, USA) as described previously (Acehan et al., 2011). Male mice were used for the experiments between 10 and 14 weeks of age. Male mice positive for the Taz shRNA transgene were identified by PCR as described previously (Acehan et al., 2011).

### MSCs isolation and culture

Mice MSCs were isolated from the femurs and tibias by flushing the bone marrow cells. The connective tissue around the bones was removed, then the bone marrow plugs were flushed using Dulbecco’s modified Eagle’s medium supplemented with 15% fetal bovine serum (FBS), 100 units/ml penicillin G, and 0.1 mg/ml streptomycin. The bone cavities were repeatedly flushed to obtain enough marrow cells. The cells were then plated and cultured in the same media at 37°C with 5% CO_2_. The medium was changed on the next day and the non-adherent cells were removed. Every 3 days the medium was replaced with fresh medium and the cells were cultured until confluency exceeded 90%. Unless otherwise indicated, all other reagents used in this study were of analytical grade and were obtained from either Thermo Fisher Scientific (Winnipeg, MB) or Sigma-Aldrich (Oakville, ON).

### B Cells isolation and co-culture of MSCs and B cells

Splenic naïve B lymphocytes were isolated from WT mice by magnetic bead negative selection using EasySep Mouse B Cell Isolation Kit (Catalog no. 19854) from STEMCELL Technologies (Vancouver, BC). B cells were cultured in RPMI 1640 media supplemented with 10% FBS, 1% antibiotic/antimycotic, and 2-mercaptoethanol at 37 °C with 5% CO_2_. Subsequent to isolation B cells were stimulated with 10 μg/ml lipopolysaccharide (LPS) (Catalog no. L2887) from Sigma-Aldrich (Oakville, ON). For co-culture with MSCs, the naïve B cells were stimulated by LPS (10 μg/ml) for 24 h then co-cultured with WT MSCs or TazKD MSCs for 72 h in 24-well plates at ratio 1:10 of MSCs:B cells.

### Metabolic assays

Oxygen consumption rate (OCR) and extracellular acidification rate (ECAR) were measured in MSCs using a Seahorse XF24 Extracellular Flux Analyzer and kits from Agilent Technologies (Santa Clara, CA). Briefly, isolated WT and TazKD MSCs were seeded in XF24 24-well plates coated at density of 4×10^5^ cells per well. For mitochondrial function stress test (OCR assay) the following final concentrations of drugs were used: 1 μM oligomycin, 1 μM FCCP, 1 μM antimycin A, and 1 μM rotenone as per the manufacturer’s instructions. Basal respiration, maximal respiration, and spare respiratory capacity were determined. For glycolytic function test (ECAR assay), 10 μM glucose, 1 μM oligomycin, and 25 μM 2-deoxyglucose were used as per the manufacturer’s instructions. The extracellular acidification rates (ECAR) for glycolysis, glycolytic capacity, and glycolytic reserve were determined. At the end of the assay, all measurements were normalized to μg of protein in each well.

### Measurement of Mitochondrial Membrane Potential

WT MSCs and TazKD MSCs were seeded onto 96-well plate until they attained 80% confluence. The mitochondrial membrane potential was assessed using the Tetramethylrhodamine, Ethyl Ester (TMRE) mitochondrial membrane potential assay kit (Catalog no. K238) from BioVision Inc. (Milpitas, CA) according to the manufacturer’s protocol. The fluorescence intensity was read at excitation=549 nm and emission=575 nm.

### MSCs and B cells proliferation assay

MSCs and B cells proliferation were measured using the CellTiter 96^®^ AQ_ueous_ cell proliferation assay kit from Promega (Madison, WI) according to manufacturer’s instructions. Cells were cultured and treated with CellTiter 96^®^ AQ_ueous_ solution and then the absorbance was recorded at 490 nm.

### Western blot analysis

Cell proteins were extracted using lysis buffer (RIPA Lysis buffer, catalog no. 89900) from Thermo Fisher (Winnipeg, MB). The total protein levels were measured by the Bradford method (Bradford, 1976). Briefly, 30 μg of protein was mixed with an equal volume of 2×Laemmli buffer supplemented with 5% 2-mercaptoethanol (Catalog no. 1610737) then loaded in 10% polyacrylamide precast gel (TGX^™^ FastCast^™^ Acrylamide Starter Kit, 10%, catalog no. 1610172) from BioRad Laboratories (Mississauga, ON). After 1 h electrophoresis, the proteins were transferred onto a polyvinylidene difluoride membrane at 4°C. This was followed by blocking the membrane using 5% milk in a solution of TBS containing 0.5% Tween 20 at room temperature for 1 h. Then, the membrane was washed with 0.5% Tween 20 three times and incubated with primary antibody an overnight at 4°C. The primary antibody was removed the next day and the membrane was incubated with secondary antibody at room temperature for 1 h. Enhanced chemiluminescence substrate (Clarity Western ECL Substrate, Catalog no.1705060) from Bio-Rad Laboratories (Mississauga, ON) was used to visualize the protein bands. The primary antibodies used in for Western blot analysis were as follows: phosphatidylinositol-3-kinase p85α (PI3K) (catalog no. sc-1637), IDO (catalog no. sc-137012), cytotoxic T lymphocyte antigen 4 (CTLA-4) (catalog no. sc-376016), IL-10 (catalog no. sc-365858), cluster of differentiation 59 (CD59) (catalog no. sc-59095) and β-Actin (catalog no. sc-47778) from Santa Cruz Biotechnology (Dallas, TX). Taz antibody was a generous gift from Dr. Steven Claypool, Johns Hopkins University. Band intensity was quantified using ImageJ software.

### B cells population doubling and % of growth rate determination

Naïve WT B cells were stimulated with 10 μg/ml LPS for 24 h, then the activated B cells were co-cultured with WT MSCs or TazKD MSCs for 72 h at a ratio of 1:10 (MSCs:B cells). After 72 h of coculture, the B cells were stained with Trypan blue and the live cells were counted. The doubling time was calculated using the following formula:

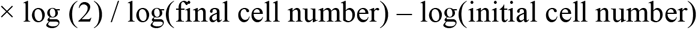

The % growth rate was calculated as follows:

Step 1: The percent change from one period to another was calculated using the following formula:

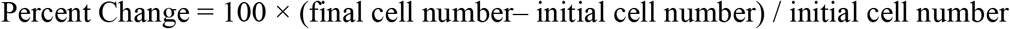

Step 2: The percent growth rate was calculated using the following formula:

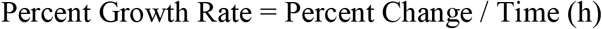

### Measurement of antibodies and IL-10 concentration

Naïve WT B cells were activated with 10 μg/ml LPS for 24 h and were then co-cultured with WT MSCs or TazKD MSCs for 72 h. The supernatant was then collected and IgG, IgM, and IL-10 levels were measured using isotype-specific enzyme-linked immunosorbent assay (ELISA) kits. IgG (total) mouse uncoated ELISA kit (Catalog no.88-50400-22) and IgM mouse uncoated ELISA kit (Catalog no.88-50470-22) were from Thermo Fisher (Winnipeg, MB). Mouse IL-10 DuoSet ELISA (Catalog no. DY417-05) was obtained from R&D Systems (Minneapolis, MN).

### Flow cytometry analysis

LPS activated B cells were co-cultured with WT MSCs or TazKD MSCs at ratio 1:10 (MSCs: B cells). The surface markers of B cells were assessed after co-culture with MSCs by flow cytometry analysis. Briefly, after 72□h of co-culture, B cells in the supernatant were collected and centrifuged at 1200 rpm for 5 min. Then, the pellet was washed 2 times with PBS and suspended in 500 μl FACS buffer (PBS containing 2% FBS). Plasma cells were analysed by staining with fluorochrome-conjugated antibodies against B220, CD19, and CD138 obtained from BioLegend (San Diego, CA). IL-10 producing B cells and Plasma cells were analysed via intracellular cytokine staining described below. The cells were stimulated for 4 h with 50 ng/mL phorbol myristic acetate 500 ng/mL ionomycin, and 10 μg/mL brefeldin A. Cells were fixed, surface stained for B220 and CD138, and subsequently stained for intracellular IL-10 (BD Biosciences, Mississauga, ON). Cells were then washed with FACS buffer then were acquired and analyzed using a BD FACS Canto II cytometer and FlowJo software (BD Biosciences), respectively. Cells were gated first on lymphocytes, single cells by forward scatter width, and then dead cells gated out using Aqua viability dye from BioLegend (San Diego, CA). Samples were then labeled with specific antibodies. Anti-CD19 (APC-Cy7), anti-B220 (PE-Cy7), anti-CD138 (APC), anti-IgD (FITC), anti-IL-10 (PE), anti-Sca1 (BV421), anti-IgM (PE-Cy7), anti-CD21 (FITC), anti-CD5 (PE) obtained from BD Biosciences. Cells were analyzed by flow cytometry using a BD FACS Canto II instrument. FlowJo software was used for data analyses.

### Lipid Mass Spectrometry

Electrospray ionization - mass spectrometry coupled to high performance liquid chromatography was performed to quantify CL mass and fatty acyl molecular species as previously described (Sparagna et al., 2005).

### Statistical analysis

All experimental results were expressed as mean ± SD. The *p* values were calculated using two tailed unpaired Student’s t test unless otherwise specified/one-way-analysis of variance followed by multiple comparisons by Tukey’s test. A p value of<□0.05 was considered statistically significant.

## Results

### Taz deficiency in MSCs increases glycolysis but does not alter mitochondrial function

Taz deficiency results in a decrease in CL (Lou et al., 2018). Initially we examined Taz expression and CL levels in MSCs. TazKD MSCs exhibited a 55% (*p*<0.0001) reduction in Taz protein expression and this was associated with a 42% (*p*=0.005) reduction in CL compared to WT MSCs (Supplementary Fig. 1A, 1B). The reduction in CL was seen across all CL molecular species examined (Supplementary Fig. 1C). We next assessed mitochondrial respiration in WT MSCs and TazKD MSCs. Surprisingly, we observed no alteration in mitochondrial OCR including basal respiration (measured before addition of inhibitors), maximal respiration (determined after addition of FCCP), or spare respiratory capacity (the difference between the maximal OCR and basal OCR) in TazKD MSCs compared to WT MSCs (Fig. 1A). Thus, mitochondrial respiration did not appear to be impacted by Taz deficiency and CL loss in MSCs. We then measured extracellular acidification rate (ECAR) in wild type MSCs and TazKD MSCs. TazKD MSCs exhibited a 45% increase (*p* =0.03) in glycolysis compared to WT MSCs (Fig. 1B). In contrast, glycolytic capacity and glycolytic reserve were not altered in TazKD MSCs (Fig. 1B). Since our metabolic analysis indicated that mitochondrial function was not altered in TazKD MSCs, as a further control, we investigated whether mitochondrial membrane potential was affected by Taz deficiency. TMRE is a positively charged red-orange fluorescent dye that accumulates in active negatively charged mitochondria. Therefore, decreased fluorescent intensity indicates the presence of inactive mitochondria. WT MSCs and TazKD MSCs were incubated with TMRE and fluorescent intensity determined. No change in fluorescent intensity was observed between WT MSCs and TazKD MSCs confirming that mitochondrial activity was unaltered in TazKD MSCs (Fig. 1C). Thus, TazKD MSCs exhibit an increase in glycolysis. In MSCs increased glycolysis is associated with a shift in their immunological properties. This interesting finding prompted us to investigate the immunomodulation abilities of TazKD MSCs.

**Figure 1.**
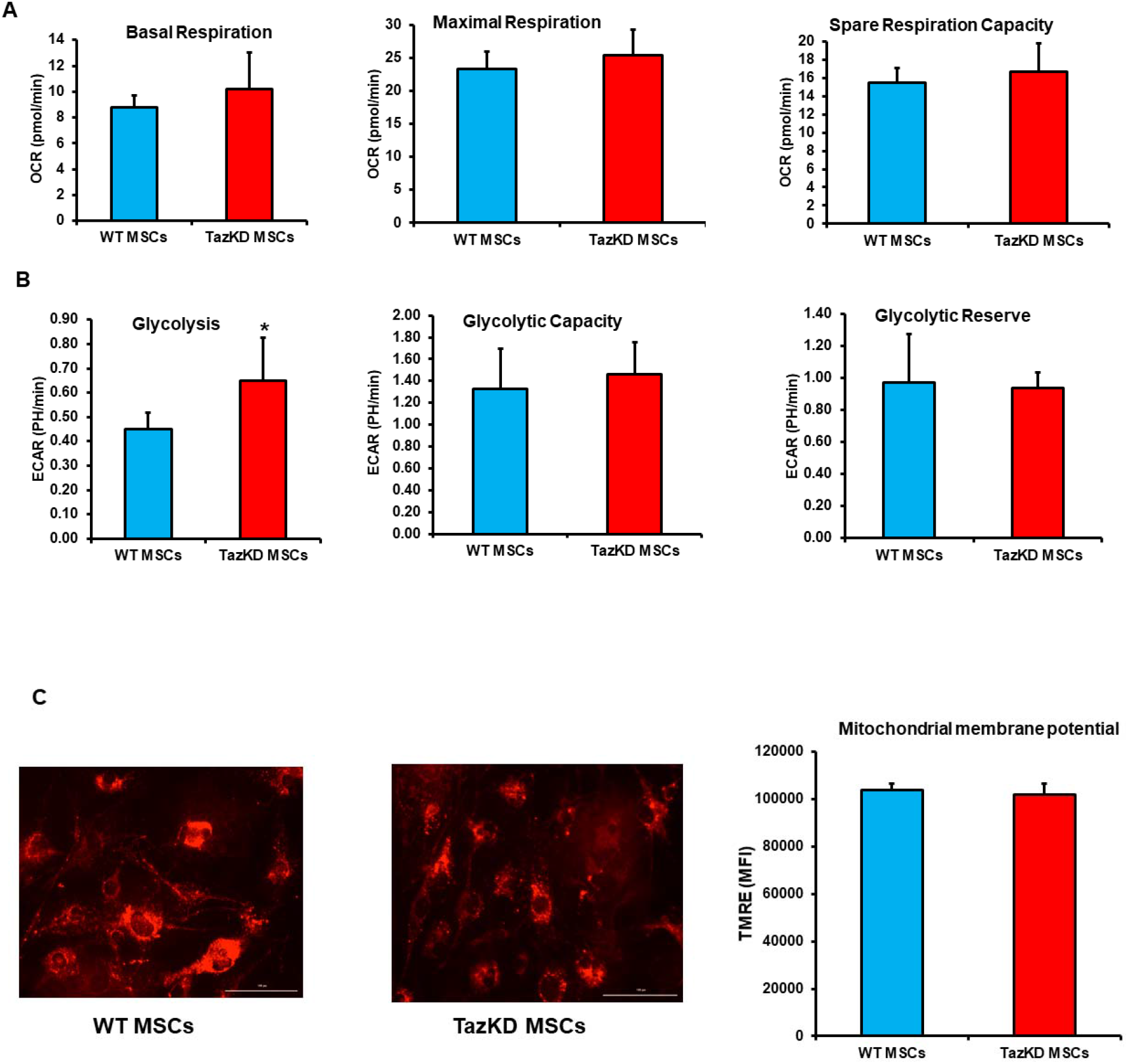
Taz deficient MSCs exhibit increased glycolysis. Mitochondrial function and glycolytic activity and mitochondrial membrane potential were determined in WT MSCs and TazKD MSCs as described in Materials and Methods. **A**. Basal respiration, maximal respiration and spare respiratory capacity. **B**. Glycolysis, glycolytic capacity and glycolytic reserve. **C.** Mitochondrial membrane potential in WT and TazKD MSCs. Results expressed as mean ± SD, n=3. \**p*<0.05 compared with WT MSCs cells.

### Tafazzin deficiency promotes MSCs proliferation and increases expression of immunosuppressive markers

Initially we examined the impact of Taz deficiency on MSCs proliferation. We found that the proliferation rate of TazKD MSCs was increased 27% (*p*=0.009) compared to WT MSCs (Fig. 2A). The PI3K pathway is essential for MSCs survival and proliferation and upregulation in PI3K expression is considered a marker for increased proliferation of MSCs (Chen et al., 2013). We found that PI3K expression was elevated 78% (*p*=0.04) in TazKD MSCs compared to WT MSCs (Fig. 2B).

**Figure 2.**
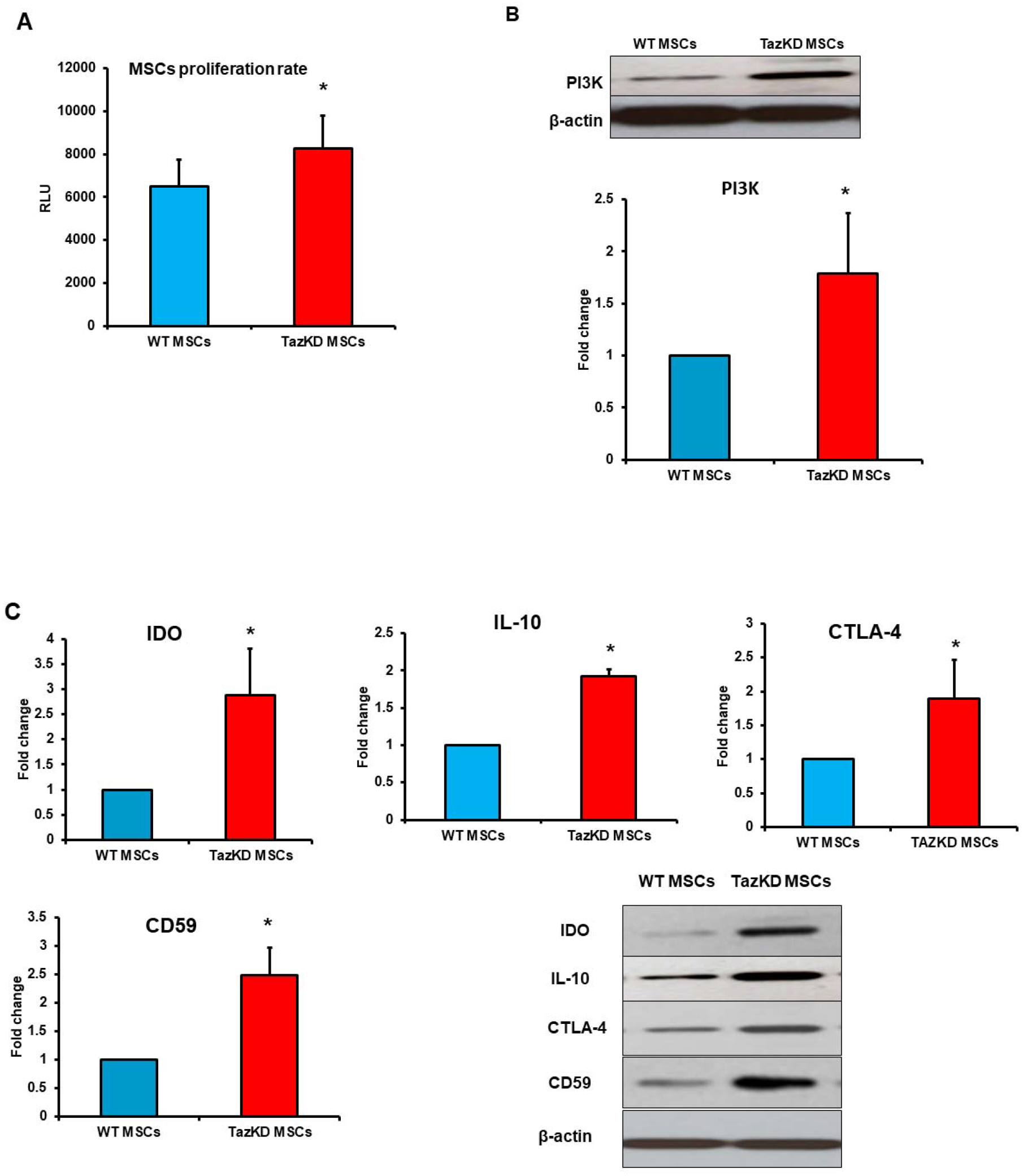
Taz deficiency promotes MSCs proliferation and increases MSCs immunosuppressive markers. Proliferation rate (**A**), PI3K protein expression (**B**), and protein levels of immunosuppressive markers IDO, IL-10, CTLA-4, CD59, and β-actin as loading control (**C**) in WT and TazKD MSCs were determined as described in Materials and Methods. Results expressed as mean ± SD, n=3. \**p*<0.05 compared with WT MSCs cells.

MSCs are immunosuppressive cells which can modulate and suppress other cells of the immune system through expression of a variety of immunosuppressant molecules including IDO, CTLA-4, IL-10, and CD59 (Ma and Chan, 2016). These markers modify the function of many immune cells including B cells and mitigate their activation (Jiang and Xu, 2020). Interestingly, protein expression of all immunosuppressive markers examined were markedly increased in TazKD MSCs compared to WT MSCs (Fig. 2C). Specifically, IDO expression was increased 187% (*p*=0.008), CTLA-4 increased 89% (*p*=0.02), IL-10 increased 192% (*p*□0.0001) and CD59 increased 148% (*p*=0.001) in TazKD MSCs compared to WT MSCs. Thus, MSCs isolated from TazKD mice exhibit a higher proliferation rate and an elevated expression of immunosuppressive markers. The increased proliferation and expression of immunosuppressive markers was consistent with the increase in glycolysis in TazKD MSCs.

### TazKD MSCs inhibit B cell proliferation and reduce the rate of B cell growth

B lymphocytes are indispensable for adaptive and humoral immunity and play a vital role in maintenance of normal immune response. Inhibition of B cell proliferation and differentiation weakens the immune system and contributes to immunodeficiency (Ahn and Cunningham-Rundles, 2009). Previous studies have shown that MSCs mitigate the proliferation of immune cells by either secreting immunosuppressive molecules or through cell-to-cell contact (Liu et al., 2020c). We thus investigated the effect of TazKD MSCs on B cell proliferation. B cells were isolated from WT mice and activated by incubation with LPS for 24 h. LPS binds to Toll-like receptor-4 and promotes murine B cell proliferation (Urbanczyk et al., 2018). Subsequently, WT MSCs or TazKD MSCs were co-cultured with activated B cells for 72 h and B cell proliferation determined. B cell proliferation was inhibited 60% (*p*=0001) when co-cultured with TazKD MSCs compared to WT MSCs (Fig. 3A). In addition, co-culture of activated B cells with TazKD MSCs increased the population doubling time of B cells by 43% (*p*= 0.003), and decreased B cell growth rate by 58% (*p*= 0.002) compared to B cells co-cultured with WT MSCs (Fig. 3B, 3C). Thus, Taz deficiency in MSCs inhibits B cell proliferation and reduces the rate of B cell growth.

**Figure 3.**
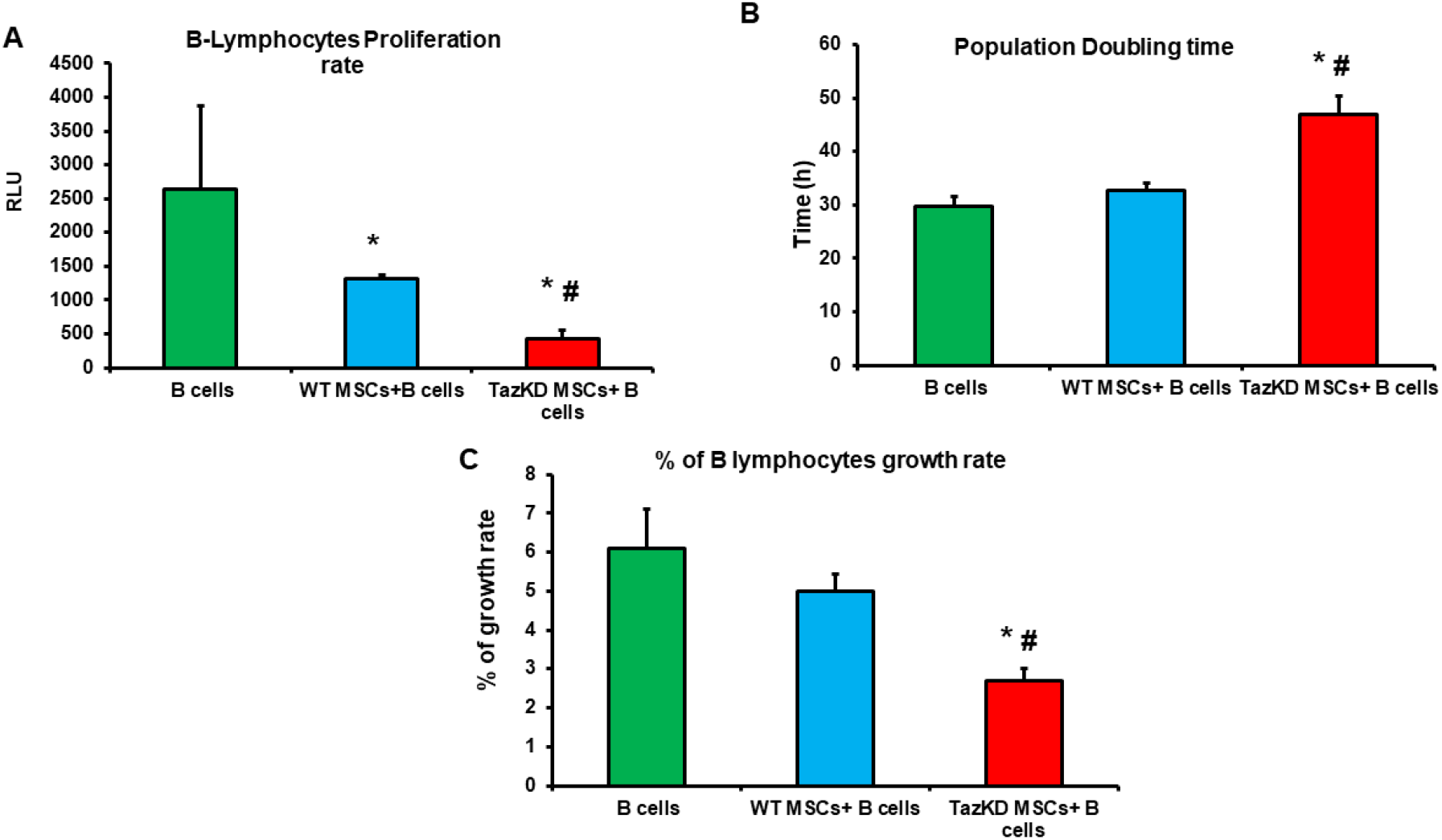
TazKD MSCs inhibit B cell proliferation and decrease B cell growth rate. WT B cells were activated with LPS then co-cultured with WT MSCs or TazKD MSCs. B cells proliferation rate (**A**), B cells population doubling time (**B**), and percent B cell growth rate (**C**) were determined as described in Materials and Methods. Results expressed as mean ± SD, n=3-5. \**p*<0.05 compared with B cells. #*p*<0.05 compared with WT MSCs.

### TazKD MSCs decrease IgM production and increase IL-10 secretion of B cells

Activation of murine B cells with LPS promote their synthesis and secretion of antibodies and MSCs have the ability to inhibit antibody secretion by LPS-stimulated B cells in co-culture (Asari et al., 2009). LPS activated B cells were co-cultured with either WT MSCs or TazKD MSCs as above and IgG and IgM secretion into the medium determined. Co-culture of B cells with TazKD MSCs reduced medium IgM levels by 46% (*p*□0.0001) compared to B cells co-cultured with WT MSCs (Fig. 4A). Medium IgG concentrations were unaltered (Fig. 4B).

**Figure 4.**
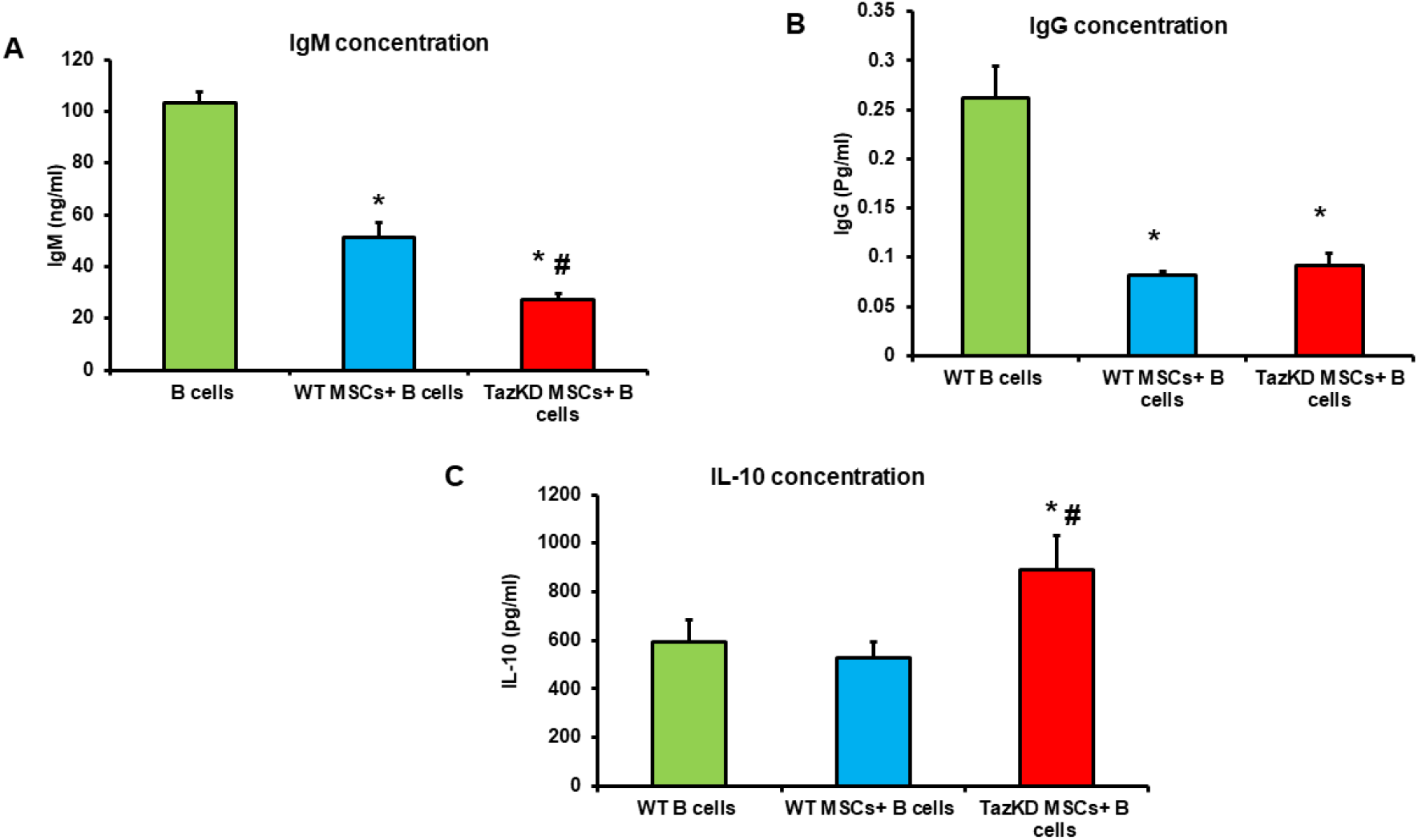
TazKD MSCs decrease B cell IgM production and increase B cell IL-10 secretion. WT B cells were activated with LPS then co-cultured with WT MSCs or TazKD MSCs and IgM (**A**), IgG (**B**), and IL-10 (**C**) secretion from B cells determined as described in Materials and Methods. Results expressed as mean ± SD, n=3-5. \**p*<0.05 compared with B cells. #*p*<0.05 compared with WT MSCs cells.

MSCs promote immune tolerance by inducing a regulatory B cells (Bregs) phenotype. Bregs suppress the immune response by secreting the anti-inflammatory cytokine IL-10 (Tarique et al., 2018). LPS activated B cells were co-cultured with either WT MSCs or TazKD MSCs as above and medium IL-10 concentration determined. Co-culture of B cells with TazKD MSCs markedly increased IL-10 levels by 70% (*p*=0.001) compared to B cells co-cultured with WT MSCs (Fig. 4C). Thus, Taz deficiency in MSCs reduces IgM production and increases IL-10 secretion of B cells.

### TazKD MSCs induce the formation of B cell immunosuppressive phenotypes

B cells differentiate into antibody secreting cells known as plasma cells. Plasma cells play a pivotal role in providing sustained and long-time immunity against many infections (Pioli, 2019). To determine if Taz deficiency impacted B cell differentiation into plasma cells, LPS activated B cells were co-cultured with either WT MSCs or TazKD MSCs as above and flow cytometry analysis for determination the presence of CD138^+^ plasma cells, IL-10 secreting plasma cells, and IL-10 secreting Bregs performed. Co-culture of B cells with TazKD MSCs increased the differentiation of B cells into plasma cells by 120% (*p*=0.001) compared to B cells co-cultured with WT MSCs (Fig. 5A). In addition, co-culture of B cells with TazKD MSCs increased the number of IL-10 secreting plasma cells by 145% (*p=*0.003) compared to B cells co-cultured with WT MSCs (Fig. 5B). Immunoglobulin D (IgD) is known to signal B cell activation and its presence on B cells indicate a mature and functional B cell that can take part in immune defense (Geisberger et al., 2006; Maity et al., 2015). Stem cell antigen 1 (Sca-1) is a known marker for hematopoietic progenitor cells and its expression is indicative of immature B cells (Fossati et al., 2010; Ichii et al., 2014). Co-culture of TazKD MSCs with B cells increased the number of Sca-1^+^/IgD^−^ B cells by 104% (*p*=0.004) compared to B cells co-cultured with WT MSCs (Fig. 5C). IL-10 secreting Bregs are a potent immunosuppressive subset of B cells. We found that co-culture of TazKD MSCs with B cells increased the population of IL-10 secreting Bregs by 45% (*p*=0.03) compared to B cells co-cultured with WT MSCs (Fig. 5D). Thus, co-culture of activated B cells with TazKD MSCs induces the formation of B cell immunosuppressive phenotypes.

**Figure 5.**
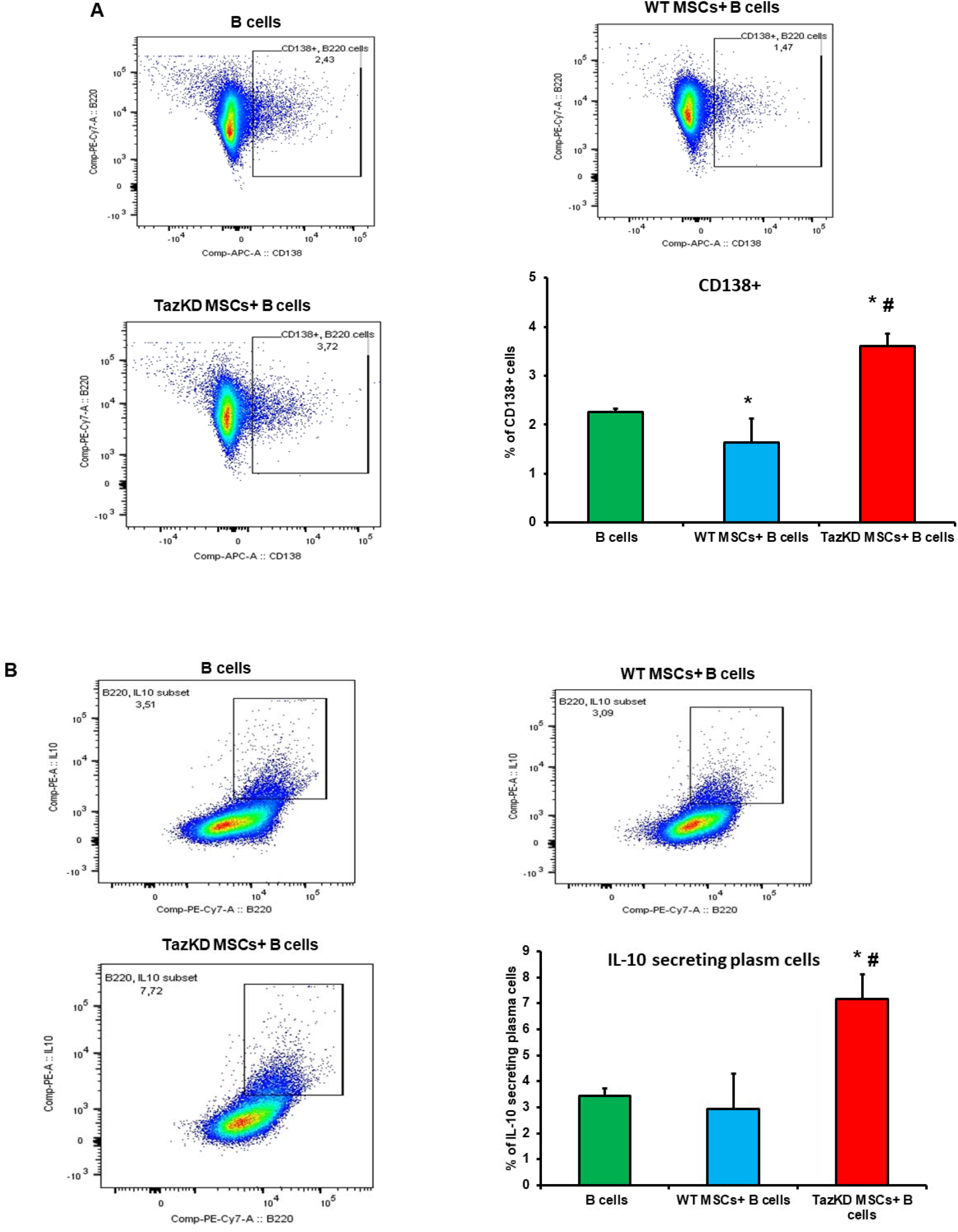
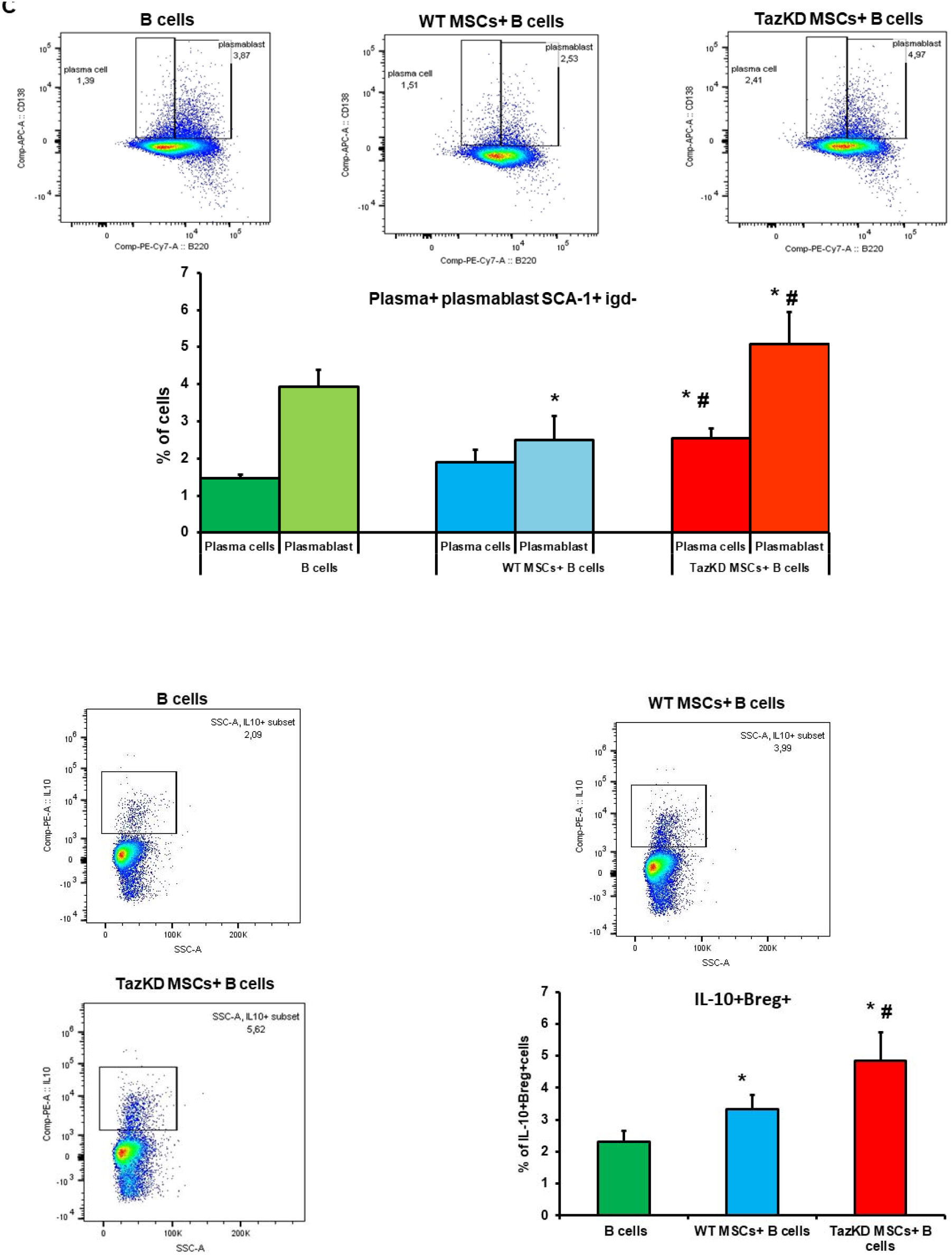
TazKD MSCs induce the formation of immunosuppressive phenotypes of B cells. WT B cells were activated with LPS then co-cultured with WT MSCs or TazKD MSCs and B cell phenotypes determined as described in Materials and Methods. Representative flow cytometry plots are shown. **A.** Percent of Plasma cells. **B.** Percent IL-10 secreting plasma cells. **C.** Percent of Sca-1^+^/IgD^−^ immature B cells. **D.** Percent of IL-10 secreting Bregs. Results expressed as mean ± SD, n=3-5. \**p*<0.05 compared with B cells. #*p*<0.05 compared with WT MSCs cells.

## Discussion

BTHS is a rare X-linked autosomal recessive disease where patients display cardiomyopathy, skeletal myopathy, growth retardation, neutropenia and susceptibility to infections. BTHS and its associated complications are caused by abnormal mitochondrial function and oxidative phosphorylation due to CL deficiency as a result of mutations in the TAFAZZIN gene which encodes the Taz protein (Saric et al., 2016). Taz is a mitochondrial inner membrane transacylase enzyme required for CL remodeling and maturation. CL is an essential lipid of the inner mitochondrial membrane and provides vital role for the proper function of many enzymes and complexes involved in oxidative phosphorylation (Paradies et al., 2019). Previous studies have reported that Taz deficiency in BTHS patients result in susceptibility to multiple and serious infections. Neutropenia has been observed in BTHS patients and this is proposed to be a link to the weaken innate immunity in these patients (Clarke et al., 2013; Dibattista et al., 2017). However, the role of Taz deficiency in other cells of the immune system is beginning to emerge. For example, Taz deficiency resulted in improper activation and differentiation of CD8^+^ T cells (Corrado et al., 2020). In addition, Taz deficiency in Mast cells impaired their exocytotic mechanisms (Maguire et al., 2021). B lymphocytes are essential for adaptive and humoral immunity. We recently reported that Taz deficiency in B cells reduced their metabolic activity and function during activation (Zegallai et al., 2021). However, the role of Taz in regulating MSCs function and the impact of Taz deficiency in MSCs on other cells of the immune system had never been investigated.

MSCs undergo a metabolic switch depending on the stress of the microenvironment or an alternation in their genetic and epigenetic characteristics (Ito and Suda, 2014). Mitochondrial membrane potential is a key determinant of mitochondrial activity and dynamics and it is maintained during normal oxidative phosphorylation (Zorova et al., 2018). Surprisingly, although Taz deficiency reduced CL in MSCs, it did not impact mitochondrial energy metabolism nor mitochondrial membrane potential. This is consistent with a previous study in which Taz knockdown did not disrupt mitochondrial morphology nor oxidative phosphorylation in hematopoietic stem and progenitor cells (Seneviratne et al., 2019).

Glycolysis is an anaerobic energy production mechanism extensively used by MSCs and increased glycolysis is associated with an increase in MSCs immunosuppressive abilities (Sigmarsdóttir et al., 2020). Remodeling of human MSCs metabolism towards glycolysis under interferon-gamma treatment potentiated stemness and immunomodulation through increased secretion of immunosuppressive markers (Liu et al., 2019). We observed that Taz deficiency in MSCs resulted in an increase in glycolysis and this was associated with increased PI3K expression indicative of increased proliferation. In addition, the increased proliferation of TazKD MSCs was associated with increased expression of the immunosuppressive markers IDO, CTLA-4, IL-10 and CD59. IDO is a soluble immunosuppressive factor that promotes B cell conversion into Bregs and thus induces an immune-tolerance state (Liu et al., 2020b). CTLA-4 is a glycoprotein belonging to the CD28 family which mediates the immune inhibitory functions of MSCs by suppressing immune-active T and B cells and induces the differentiation of these cells toward inhibitory phenotypes (Gaber et al., 2018). Moreover, CTLA-4 induces the secretion of IDO from MSCs. In addition, CTLA-4 expression potentiates the PI3K signalling pathway in MSCs (Oyewole-Said et al., 2020, Schneider et al., 2008). IL-10 is a cytokine that has many immunosuppressive effects on B cells including promoting formation of a Bregs phenotype and promoting plasma cells secretion of IL-10 (Matsumoto et al., 2014). Complement system activation is required to stimulate the migration and maturation of B cells. MSCs express CD59 which suppress activation of complement system and as a result eliminate complement system activation of B cells (Soland et al., 2013). The observed elevation in IDO, IL-10, CTLA-4 and CD59 in TazKD MSCs implicate Taz in the regulation of the immunomodulatory functions of MSCs and indicate that Taz deficiency in MSCs may result in the adoption of a more immunosuppressive state.

MSCs are known to induce an immune tolerance state, abrogate activation of B cells and dictate B cell differentiation fate. MSC’s inherited immunosuppressive abilities may diminish the proliferation and activation of B cells and switch their phenotype toward potent immune-suppressive cells including Bregs and IL-10 secreting plasma cells (Cho et al., 2017, Rosado et al., 2015). Excessive accumulation of plasma cells is associated with the development of many autoimmune diseases and allograft rejection, while under-stimulation of these cells produces weaken short-term immunity (Hoyer et al., 2004; Moise et al., 2010). We observed that the plasma cells induced by TazKD MSCs were a specialized subset known as IL-10 secreting plasma cells which exhibit a powerful anti-inflammatory ability and produce an immune tolerance state that subdues the immune response. IL-10 secreting Bregs play a central role in maintaining immunological homeostasis by skewing T cells differentiation toward potent immunosuppressive T regulatory cells and inhibit many pro-inflammatory immune cells (Dai et al., 2017). The percentage of IL-10 secreting Bregs, IL-10 producing plasma cells and Sca-1^+^/IgD immature B cells were increased by co-culture of TazKD MSCs with B cells. Thus, TazKD MSCs induced a shift in B cell differentiation towards a more highly immunosuppressive phenotype.

In conclusion, our study has demonstrated for the first time that Taz deficiency in MSCs results in modulation of their immunosuppressive activity through increased proliferation and expression of immunosuppressive markers. In addition, co-culture of Taz deficient MSCs with LPS activated WT B cells significantly inhibits B cell proliferation, decreases B cell IgM secretion, increases B cell IL-10 production and alters B cell phenotype by inducing the formation of immunosuppressive IL-10 secreting plasma cells and IL-10 secreting Bregs. Thus, it is possible that Taz deficiency in MSCs may additionally contribute to immune dysfunction in BTHS patients. The above observations may contribute to the development of new therapeutic approaches to treat infections in BTHS.

## Acknowledgements

We thank Marilyne Vandel for technical assistance. H.Z. is the recipient of a studentship from the Libyan North American Scholarship Program. L.K.C. was the recipient of an IMPACT Heart and Stroke Foundation of Canada and Canadian Institutes of Health Research Fellowships. This work was supported by Heart and Stroke Foundation of Canada, Natural Sciences and Engineering Research Council of Canada, Children’s Hospital Research Institute of Manitoba, the University of Manitoba Research Grants Program (to G.M.H.) and the National Institutes of Health (NIH) USA (P30DK048520). G.M.H. is the Canada Research Chair in Molecular Cardiolipin Metabolism.

## Author contributions

H.M.Z., E.A.-E.-R., and G.M.H. were responsible for conceptual design of the study. H.M.Z, E.A.-E.R., G.C.S. performed the experiments and analyzed the data. F.O.-A. and A.J.M. designed, performed and analyzed the Flow cytometry experiments. L.K.C. was responsible of establishing and maintaining the Tazknockdown transgenic mouse model and culling the animals. H.M.Z., E.A.-E.-R., and G.M.H. interpreted the data, carried out statistical analysis and wrote the manuscript. All authors read and approved the manuscript.

## Supplementary Figure Legend

**Supplementary Figure 1.**
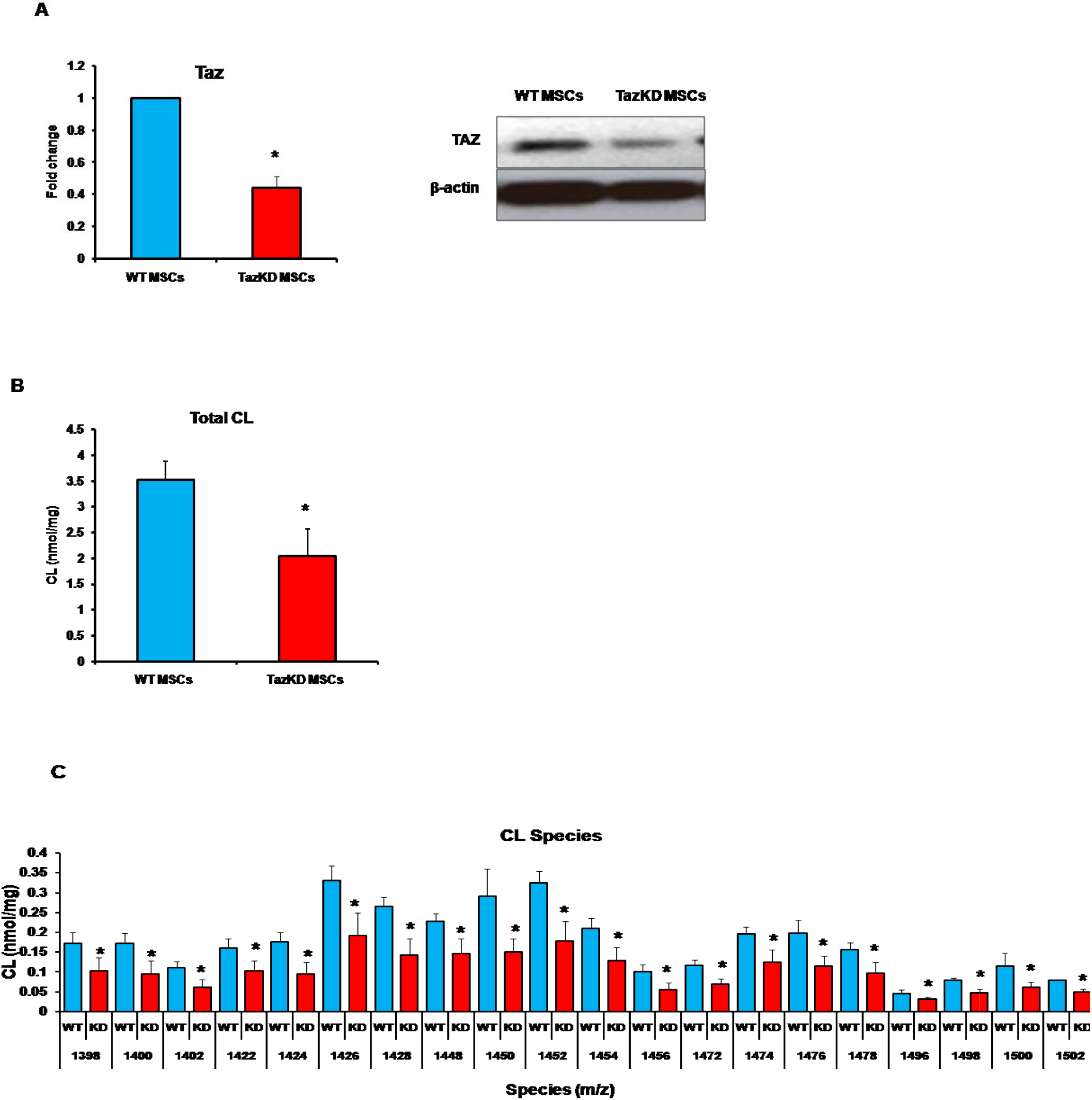
Tafazzin deficiency reduces total CL and fatty acyl molecular species in TazKD MSCs. A. Taz protein expression in WT and TazKD MSCs. B. Total CL in WT and TazKD MSCs. C. Fatty acyl species CL in WT and TazKD MSCs. n=3. \**p*<0.05 compared with WT MSCs cells.

